# Stimulation of the posterior cingulate impairs episodic memory encoding

**DOI:** 10.1101/497818

**Authors:** Vaidehi S. Natu, Jui-Jui Lin, Alexis Burks, Akshay Arora, Michael D. Rugg, Bradley Lega

## Abstract

Neuroimaging experiments implicate the posterior cingulate cortex (PCC) in episodic memory processing, making it a potential target for responsive neuromodulation strategies outside of the hippocampal network. However, causal evidence for the role PCC plays in memory encoding is lacking. In patients undergoing seizure mapping, we investigated functional properties of the PCC using deep brain stimulation (DBS) and stereotactic electroencephalography (stereo EEG). These techniques allow precise targeting of deep cortical structures including the PCC, and simultaneous acquisition of oscillatory recordings from neighboring regions such as the hippocampus. We used a free recall experiment in which PCC was stimulated during item encoding period of half of the study lists, while no stimulation was applied during encoding period of the remaining lists. We evaluated if stimulation affected memory and/or modulated hippocampal activity. Results revealed four main findings. (i) Stimulation during encoding impaired memory for early items on the study lists. (ii) Stimulation increased hippocampal gamma band power. (iii) Stimulation-induced gamma power predicted memory impairment. (iv) Functional connectivity between the hippocampus and PCC predicted the degree of stimulation effect on memory. Our findings offer the first causal evidence implicating the PCC in episodic memory encoding. Importantly, results highlight that stimulation targeted outside of the temporal lobe can modulate hippocampal activity with implications on behavior. Furthermore, *a-priori* measures of connectivity between brain regions within a functional network can be informative in predicting behavioral effects of stimulation. Our findings have significant implications for developing therapies to treat diseases of memory loss and cognitive impairment using DBS.

**Significance Statement:** Cognitive impairment and memory loss are critical public health challenges. Deep brain stimulation (DBS) is a promising tool for developing strategies to ameliorate memory disorders by targeting brain regions involved in mnemonic processing. Using DBS, our study sheds light on the lesser-known role of the posterior cingulate cortex (PCC) in memory encoding. Stimulating the PCC during encoding impairs subsequent recall memory. The degree of impairment is predictedby stimulation-induced hippocampal gamma oscillations and functional connectivity between PCC and hippocampus. Our findings provide the first causal evidence implicating PCC in memory encoding and highlight the PCC as a favorable target for neuromodulation strategies, using *a-priori* connectivity measures to predict stimulation effects. This has significant implications for developing therapies for memory diseases.

## Introduction

Deep brain stimulation (DBS) is a powerful tool used clinically to target dysregulated neural circuits and to treat a range of neurological disorders including Parkinson’s disease, essential tremor, psychiatric illnesses, and epilepsy (1–6). Recently, a large number of studies have applied DBS in patients undergoing treatments for epilepsy for probing core regions involved in memory processing, including the hippocampus, entorhinal cortex, and lateral temporal lobe and for understanding the effects of stimulation on memory performance (7–14). Results predominantly show that applying electrical stimulation to the hippocampus and entorhinal cortex impairs memory (11, 15) (with some exceptions (10)), providing evidence for causal role of these regions in mnemonic processing. This makes DBS ideal for studying cognitive enhancement and causal roles of deep cortical structures associated with memory (7–14, 16) and sensory processes (17, 18) as well as for developing neuromodulation strategies to treat memory and motors disorders (19).

One deep brain region associated with episodic memory processing is the posterior cingulate cortex (PCC). Although the PCC is not a core memory region like the hippocampus, non-invasive functional magnetic resonance imaging (fMRI) studies consistently find reduced neural activation for remembered items than forgotten items (a negative subsequent memory effect, SME) during memory encoding in the PCC (20–24). Subsequent memory effects selectively associated with encoding of spatial location and the color of stimuli have also been reported in the PCC (25). By contrast, the PCC exhibits increased activation during retrieval in fMRI investigations (26–30) and the preferential role of the PCC during memory retrieval is commensurate with evidence of its participation in the default mode network (31, 32). Furthermore, anatomical connections between PCC and temporal and frontal brain regions via the cingulum bundle, and clinical memory impairments associated with reduced metabolism in PCC and abnormal functional connectivity between PCC and hippocampus (33–38), implicate that the PCC may be a part of the extended hippocampal memory system (39). Based on this prior evidence, we sought to use DBS to test (i) the causal role of PCC in episodic memory encoding and (ii) if PCC could be used as a neuromodulation target outside of the hippocampal network for memory enhancement.

We reasoned that if neural activation in the PCC predominantly reduces during successful memory encoding (20–24), then high frequency stimulation delivered to the PCC during encoding might enhance memory. To test our hypothesis, we applied stimulation using robotically placed stereotactic electroencephalography (stereo EEG) electrodes implanted for seizure mapping, based on our prior work (40). This unique technique allows: (i) precise targeting of deep brain regions such as the PCC (**Fig. 1a**) and (ii) simultaneous acquisition of oscillatory recordings from neighboring cortical regions such as the hippocampus (**Fig. 1b**). We designed a free recall task (**Fig. 2a** and Methods) in which participants (***N***=17) first memorized words from a study list in an encoding period, which was followed by a 20s mathematical distractor task, following which participants were asked to freely recall words from memory during a retrieval phase. PCC was stimulated during the entire encoding period of half of the total number of study lists, to effectively test for prolonged list-level effects. No stimulation was applied during the encoding period of the remaining half of the study lists. Across all participants, we tested if: (i) stimulation of PCC induced memory enhancement or impairment, (ii) stimulation of PCC induced modulations in hippocampal oscillations during successful and unsuccessful memory encoding and (iii) there was a relationship between behavioral and neural effects of stimulation.

**Figure 1.**
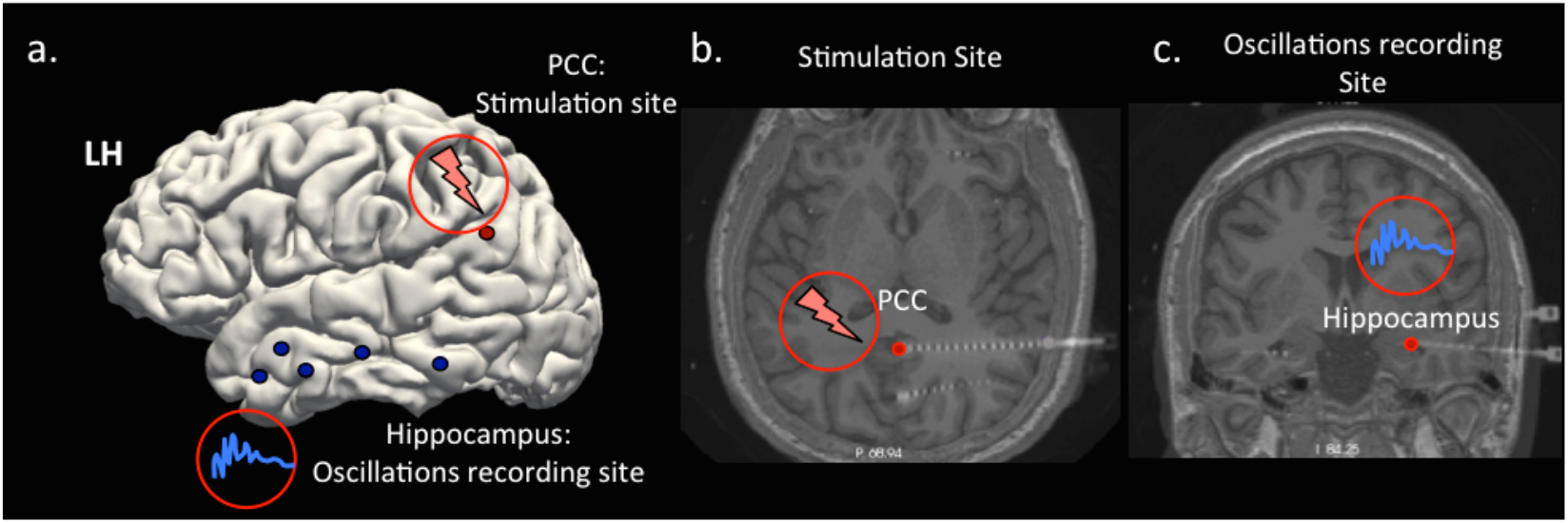
Electrode placement during deep brain stimulation (DBS) paradigm in a sample participant. a) Schematic represents a Freesurfer-based (41) pial surface in a sample participant, showing the location of electrodes in (i) the posterior cingulate cortex (PCC, red dot), where we applied electrical stimulation and (ii) the hippocampus (blue dots) where we measured oscillatory recordings. b-c) Transverse and coronal slices of the sample brain showing the deep insertion of stereo electrodes in the PCC and hippocampus, respectively, in the left hemisphere. DBS was applied to the PCC during the encoding phase in a free recall task and neural oscillations were recorded from the hippocampus.

**Figure 2.**
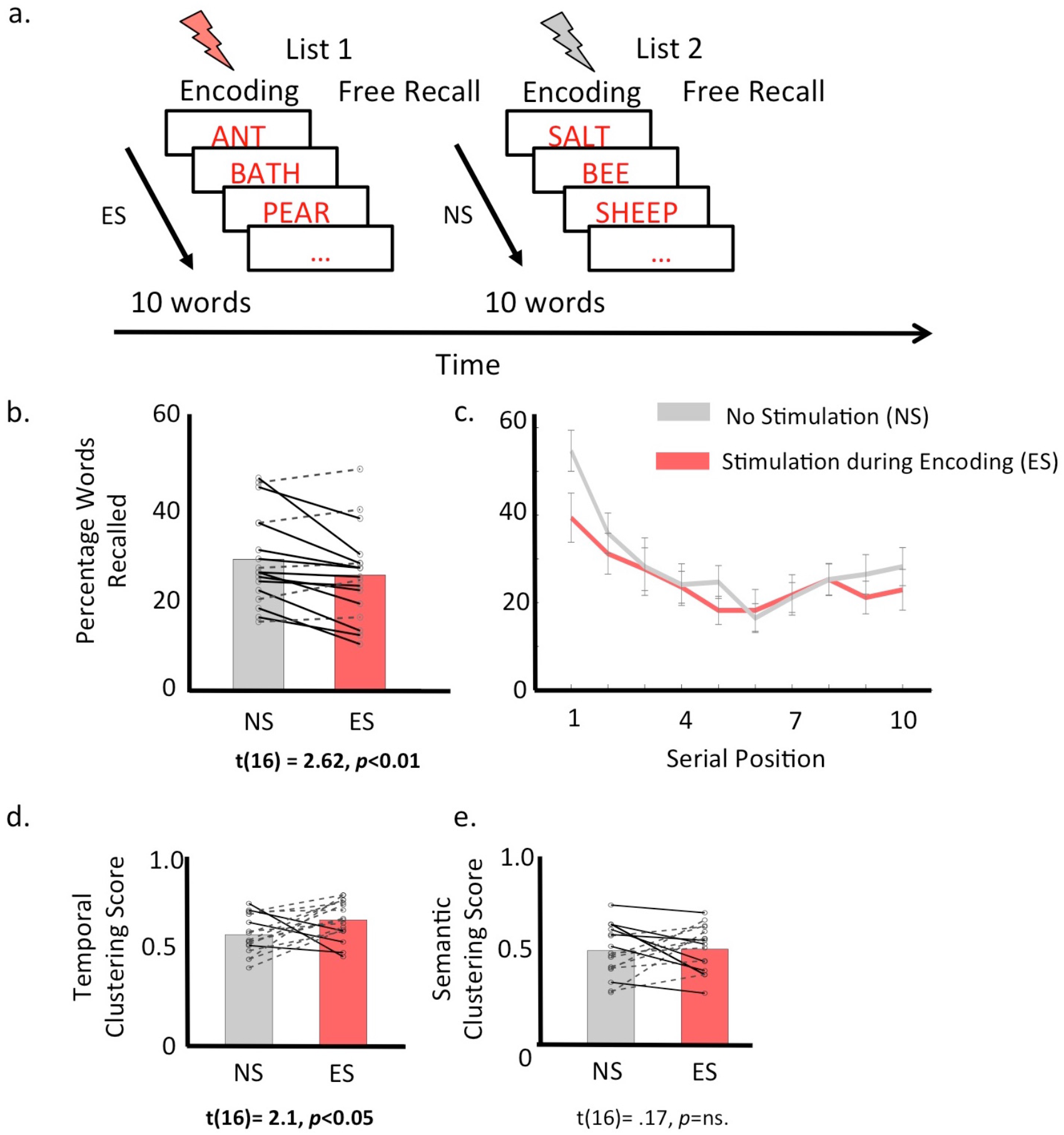
Stimulating PCC during item encoding disrupts verbal recall memory. a) A schematic of the free recall paradigm showing alternate blocks of study lists, in which participants either received stimulation during the entire word encoding period (ES, orange lightening bar) or they received no stimulation (NS, gray lightening bar). Participants learned 10 words per list and were asked to recall words from memory following a 20 s mathematical distractor task. b) Participants correctly recalled less number of words when PCC was stimulated during encoding than when no stimulation was applied. c) Serial position curves showing percentage of correctly recalled words as a function of item position during memory encoding, in NS (gray line) versus ES (orange line) conditions. Primacy effect i.e., the percentage of correctly recalling early than late items in a list was also lower in ES than NS condition. d) In contrast, temporal clustering score (TCF) was slightly higher in the ES than NS condition, whereas e.) semantic clustering score (SCF) was comparable across the two conditions. In figures b, d, and e: solid lines represent participants that showed a reduction in the measured quantity with stimulation, dotted lines represent participants that showed an increase in measured quantity with stimulation.

## Results

### Stimulation of PCC impairs verbal recall memory

First, to determine if stimulating the PCC enhances memory, we used a free recall memory paradigm in which PCC was stimulated during memory encoding period of half of the study lists (**Fig. 2a**). We compared recall performance, measured as the percentage of correctly recalled study words, when stimulation was applied during encoding period of a study list (i.e. stimulation during encoding (ES) condition) versus when stimulation was not applied (i.e. no stimulation (NS) condition). Data from all participants (N=17) were included in the analysis as all participants recalled more than 10% of the words in the experiment. Stimulation did not elicit any reports of subjective experiences, and nor was there evidence of induced seizures or other adverse consequences of high frequency 100 Hz, 2mAmps stimulation. Results revealed that PCC stimulation during encoding significantly disrupted recall performance (M_NS_ = 28.9±10.2, M_ES_ = 25.3±10.2, t(16) = 2.62, *p*<0.01, **Fig. 2b**). This performance reduction was not a consequence of list intrusions, that is the number of intrusions measured as the recall of words from prior lists, as the number of intrusions were comparable across NS and ES conditions (M_NS_ = 4.17±3.39, M_ES_ = 4.52±3.62, t(16) = 0.49, *p*=ns).

Next, we asked if stimulation-related recall impairment demonstrated a serial order effect. Specifically, we examined performance separately for primacy items (items in the 1st position in each 10-word list) and non-primacy items (items in 2nd to 10^th^ positions in each list). Serial position curves are shown in **Fig. 2c**. First, in line with long-standing findings (42), recall probability for primacy items was higher than that for non-primacy items (main effect of item position in a 2-way analysis of variance (ANOVA) with factors item position (primacy/non-primacy items) and stimulation condition (ES/NS), F_1,336_=49.24, *p*<0.001). Importantly, PCC stimulation affected recall of primacy items to a greater extent than recall of nonprimacy items (item position x stimulation condition: F_1,336_=4.08, *p*<0.05). Specifically, recall memory for primacy items in ES lists was lower than NS lists (t(16)=3.98, *p*<0.001), but was comparable for nonprimacy items (t(16)=1.46, *p*=n.s.).

In a free recall task participants tend to remember items in the order that they were encoded (43). Participants also have a tendency to cluster recall items based on their semantic association. Thus, next we asked if stimulating the PCC disrupted temporal and/or semantic organization of memory, an effect driven by hippocampal-dependent associative encoding (44). We measured the temporal clustering factor (TCF (45, 46), Methods), which is a measure of subject’s tendency to cluster the recalled items based on their temporal proximity during the study period. In contrast to recall performance, which decreased when PCC was stimulated, TCF was slightly *greater* in ES than NS conditions (t(16) = 2.13, *p*<0.05, **Fig. 2d**). This indicates that while stimulation disrupts primacy, it perhaps disposes memory circuits towards temporal associative encoding. Next, we also measured the semantic clustering factor (SCF (45, 46), Methods), a distance measure of participants’ tendency to cluster recall items by semantic associations. Results showed comparable SCF in ES and NS lists (t(16) = 0.18, *p*=ns, **Fig. 2e**). Thus, in contrast to recall performance and primacy that decreased when PCC was stimulated, associative memory processes including the TCF and SCF remain largely unaffected. This result is different than that observed when stimulating core memory regions including the entorhinal cortex and hippocampus as stimulation in these sites disrupts all aspects of memory processing including temporal clustering (11, 15).

### Stimulation of PCC increases gamma-band and decreases theta-band activation in the hippocampus

Next, using the unique opportunity afforded by the stereotactic EEG implantation technique that allowed simultaneous recording in the hippocampus while stimulating the PCC, we examined if stimulation modulates neural activity in hippocampus. For each subject (across electrodes, time windows and frequencies) hippocampal power was extracted per study item (including successfully remembered and forgotten items) following item presentation for ES and NS conditions. We then contrasted power across the two conditions using t-statistics (Methods). Across all participants, the distribution of t-statistics was compared to zero using an unpaired t-test, resulting in *p*-values, converted to z-scores, for each time window and frequency (significant frequency band clusters across participants were obtained at *p*<0.05, corrected for multiple comparisons). Results revealed that stimulation to PCC altered hippocampal activity. **Figure. 3a** shows the direction of power changes in a sample participant across low and high frequency bands. Positive t-values correspond to an increase in power in ES versus NS conditions. Specifically, we found that hippocampal power decreased in theta band (2-10 Hz) but increased in low gamma (25-40 Hz) and high gamma bands (>40Hz) in the ES compared to NS condition across all participants (**Fig. 3b**). Thus, results indicated that stimulating the PCC during memory encoding impacts hippocampal oscillations.

**Figure 3.**
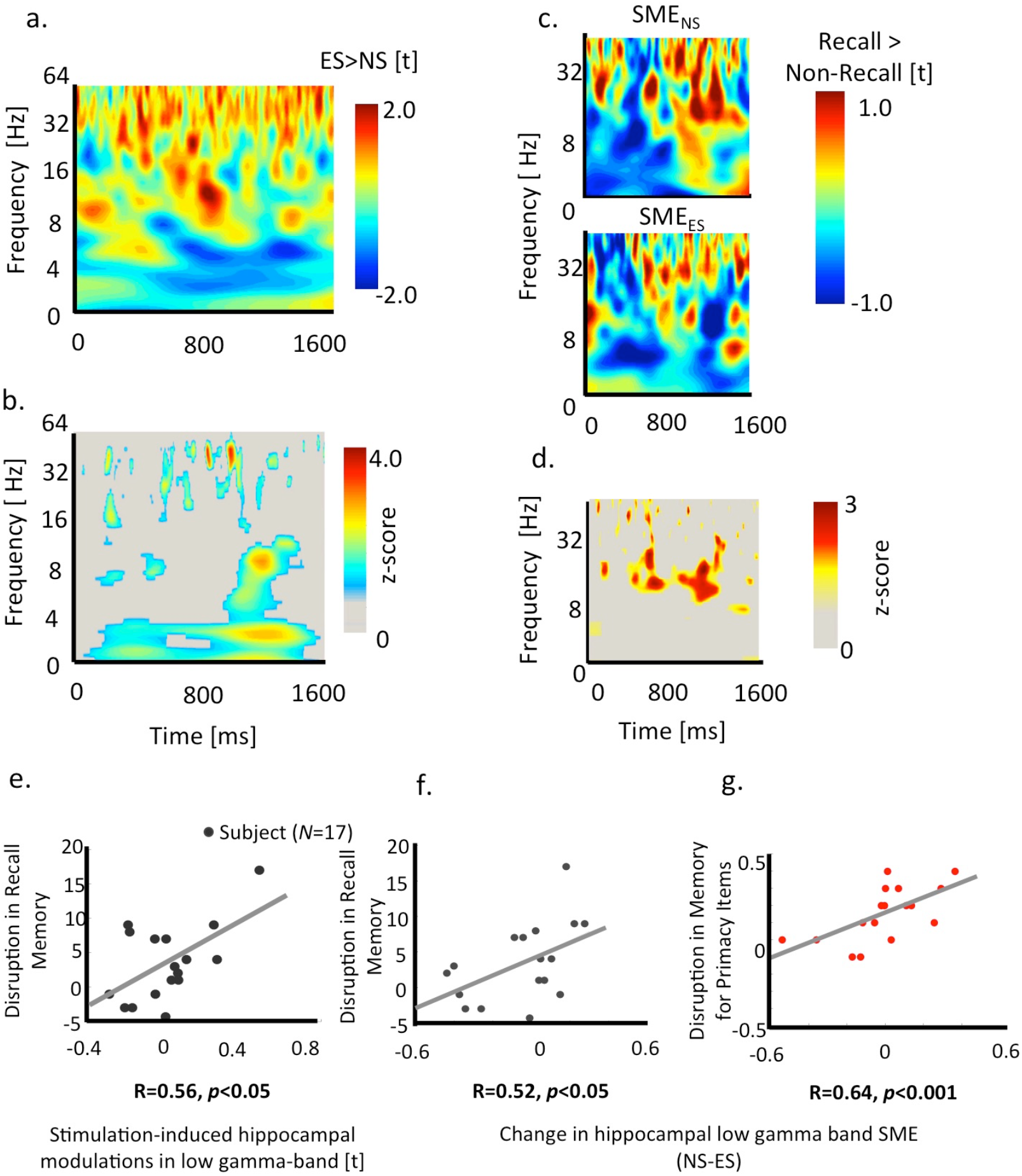
Stimulation induces changes in low gamma and theta band oscillations in the hippocampus and neural changes are linked to behavioral memory disruptions. a.) Frequency-time plot of a sample subject showing increases in low and high gamma band power, and decrease in theta band power during ES versus NS conditions. b.) Clusters of theta and gamma band power showing significant stimulation-induced power changes across all participants. (c) Frequency-time plot of a sample subject showing power differences when an item is successfully recalled versus forgotten (successful memory encoding, SME) in ES (top) and NS (bottom) conditions. Gamma power increased where as theta power reduced in both cases. (d) Clusters of theta, beta, and gamma band power showing significant stimulation-induced power changes in delta SME (SME_NS_-SME_ES_) across all participants. (e) Significant positive correlation between stimulation-induced hippocampal gamma power changes (measured as t-statistics of the contrast between ES versus NS conditions) and disruption in the memory (measured as percent recall during NS minus ES conditions). (f-g) Significant positive correlation between delta hippocampal gamma SME (measured as: SME in NS minus SME in ES condition) and disruption in memory performance (measured as: percent recall of all items/percent recall of primacy items in NS minus that in ES condition).

Next, we examined how hippocampal oscillatory power differed between successfully remembered items and forgotten items (also known as hippocampal subsequent memory effect (SME (47)), in ES versus NS conditions. Electrophysiological studies show that SMEs are generally characterized by increase in gamma band power and decrease in theta band power across a broad range of brain regions, including the hippocampus (44, 48, 49). Thus, for each participant, we measured SMEs for NS and ES conditions separately (Methods). First, as noted in prior work (44), we observed an increase in gamma band and decrease in theta band power in both NS and ES conditions for successfully remembered items versus forgotten items (**Fig. 3c**). However, evaluating how SME oscillations differ between NS versus ES conditions (measured as ΔSME=SME_ns_-SME_ES_), results showed stimulation-induced reduction in power in low gamma (25-40Hz), high gamma (>40Hz), and beta (15-24 Hz) bands, but an increase in power in theta band (significant frequency band clusters across participants at *p*<0.05, corrected, **Fig. 3d**). Notably, stimulation-induced reduction in gamma band power during successful memory encoding is contrary to the overall increase in gamma band activation during stimulation observed in the previous result. We speculate that stimulating the PCC enhances overall hippocampal gamma band power for remembered as well as forgotten items, thus inhibiting normal oscillatory processes that distinguish between the two memory traces when stimulation is absent.

### Are neural modulations in gamma and theta activity related to memory performance?

Thus far results show that PCC stimulation disrupts behavioral memory performance and differentially modulates hippocampal activity, predominantly in theta, low gamma, and high gamma frequency bands. Thus, next we asked if there is a relationship between stimulation based memory disruption and hippocampal power changes. To address this question, we first correlated measures of hippocampal neural modulation due to stimulation with the disruptive effects of stimulation on recall performance. For each subject, we operationalized stimulation-induced neural changes as contrasts of power estimates for ES versus NS conditions, across all frequencies in (i) theta (2-10 Hz), low (25-40 Hz) and high gamma bands (>40Hz), then averaged t-statistics across frequencies within each frequency band. We measured disruption in memory performance as the difference in recall performance for all items as well as for only primacy items during NS versus ES conditions. Greater positive values correspond to larger disruption of memory performance due to stimulation. Results revealed that there was a significant positive relationship between behavioral and neural effects of stimulation (R=0.56, *p*<0.05, **Fig. 3e**). Specifically, participants who showed greater stimulation-induced disruption in recall memory also showed a larger increase in power in low gamma band. A similar relationship between behavioral and neural effects of stimulation was not evident for theta and high gamma bands or for stimulation-based primacy effect (Rs<0.32, *ps*=ns).

Next, we tested if stimulation-induced changes in recall performance for all and primacy items were linked to SME related neural modulations in theta, beta, and gamma band SMEs. For each subject, we first averaged t-statistics contrasts between successful versus unsuccessful items encoding across all frequencies in (i) theta, beta, low and high gamma bands for NS and ES conditions separately, then we subtracted averaged power NS and ES conditions for each frequency bands and correlated with behavioral differences in recall performance for all items and only primacy items during NS and ES conditions. Results revealed a significant positive relationship between stimulation-induced memory disruption for all items and SME related power changes in low gamma band (R=0.52, *p*<0.05, **Fig. 3f**). A similar significant relationship was found between disruption in recall memory for primacy items and power change in low gamma (R=0.64, *p*<0.001) and beta (R=0.70, *p*<0.001) bands. No significant relationship between neural and behavioral effects of stimulation was evident in theta or high gamma bands SMEs (Rs<0.04, *ps*=ns). Combined, our results indicate that PCC stimulation during encoding results in differential changes in the hippocampal neural activity, and that stimulation-induced hippocampal modulations predominantly in the low gamma band predict behavioral memory disruptions.

### Are stimulation effects on behavior and neural responses linked to functional connectivity between PCC and hippocampus?

One of the factors that can drive stimulation effects is the connectivity between brains regions (50–52). Not only is PCC anatomically connected to the hippocampus, via the cingulum bundle, but prior research also shows that abnormal functional connectivity between PCC and hippocampus may lead to cognitive impairments (38). Thus, we asked if functional connectivity between PCC and hippocampus would predict the degree of stimulation effects on recall memory and/or oscillatory changes in hippocampal activity. To exclude any stimulation related artifacts and to obtain robust connectivity measurements across a larger number of trials, we sought to measure *baseline* functional connectivity when no stimulation was present during the entire experiment (**Fig. 4a**). A subset of our participants (11 out of 17 participants) performed a follow-up free recall task without the presence of any stimulation. We obtained baseline functional connectivity measures using wavelet-extracted power and phase locking values as oscillatory synchrony between the hippocampal and PCC electrodes (Methods). We then correlated averaged connectivity phase locking statistics (53) in theta (2-10Hz), alpha (11-14Hz), beta, (15-24Hz), low (25-40Hz), and high gamma (>40 Hz) bands with stimulation-induced changes in (i) recall performance, (ii) low gamma oscillations, and (iii) delta SMEs. Interestingly, only baseline connectivity in the low gamma band was related to stimulation-induced changes in memory and neural responses. Particularly, participants with larger hippocampal-PCC connectivity in low gamma band showed greater degree of recall memory disruption due to stimulation (R=0.62, *p*<0.05, **Fig. 4b**). There were also trends for positive relationships between low gamma-band connectivity and stimulation-induced low gamma power differences (R=0.42, *p*=0.1) and SME related low gamma power changes (R=0.52, *p*=0.08). Combined, our results showed that baseline hippocampal-PCC connectivity is related to the effect of PCC stimulation on episodic memory recall and hippocampal activations. This suggests that functional connectivity may serve as *a-priori* information to predict stimulation-induced effects using DBS.

**Figure 4.**
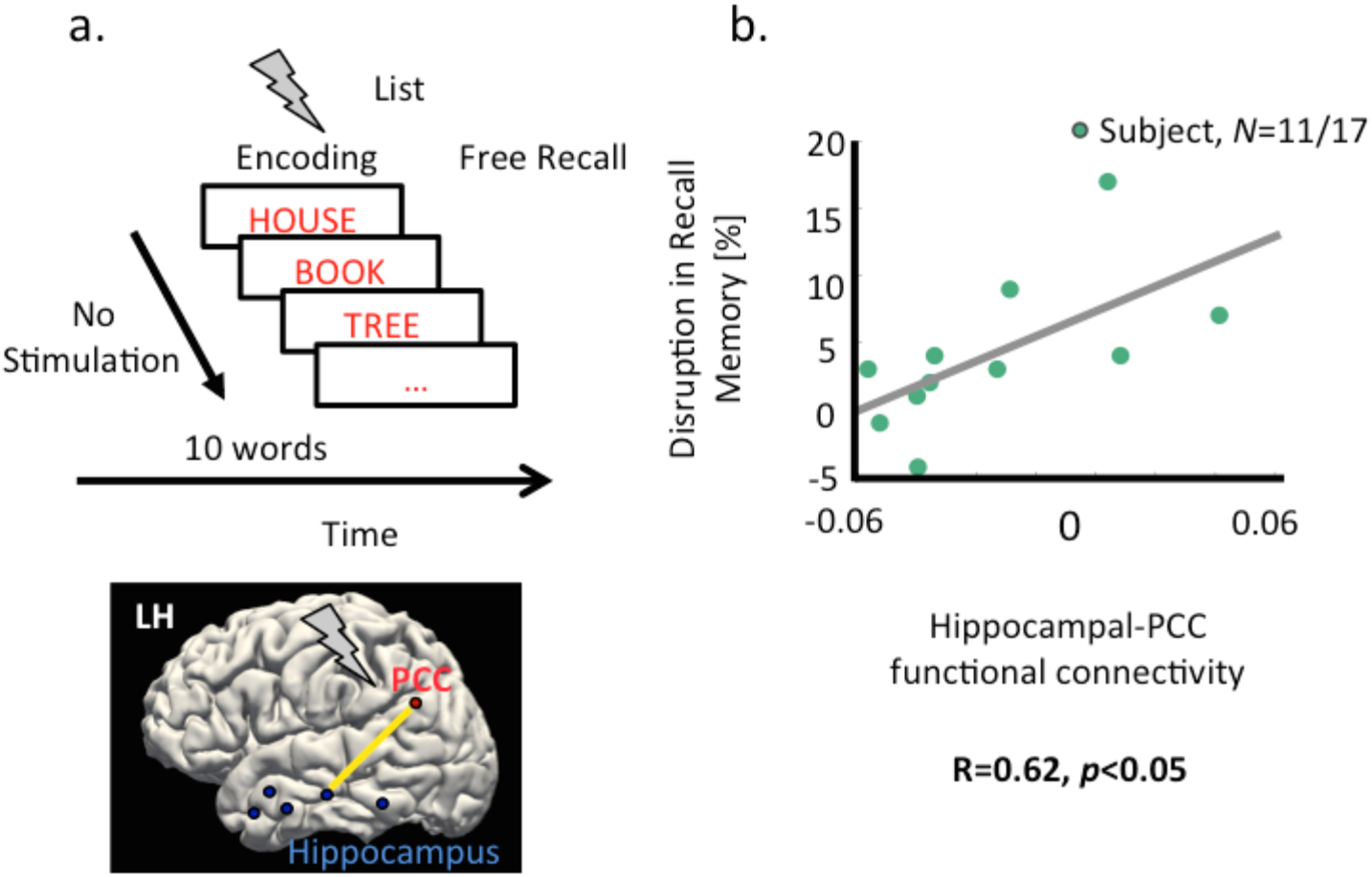
Baseline PCC-hippocampal functional connectivity is positively linked to stimulation-induced memory disruptions. a) PCC-Hippocampal functional connectivity, measured as oscillatory synchrony between hippocampal electrodes and PCC electrodes (in a subset of participants, *N*: 11 out of 17) in a second free recall experiment without the presence of stimulation. b) Positive correlation between baseline functional connectivity and stimulation-induced recall memory disruption. That is, participants with higher hippocampal-PCC connectivity tended to show larger disruption in memory performance.

## Discussion

For the first time, using DBS, our study examined the causal role of the PCC in episodic memory encoding. Our results revealed that stimulating the PCC during memory encoding disrupts, and not enhances, memory performance. Results also showed that stimulating the PCC induces modulations in hippocampal oscillatory power, with an overall increase in low gamma band power and a decrease in theta band power. However, during successful memory encoding (that is comparing power changes between successfully remembered versus forgotten items (44, 47)), stimulation resulted in decreased gamma band SME. Importantly, these neural modulations in the hippocampus are linked to memory disruptions during encoding. We also find that baseline functional connectivity between the PCC and hippocampus predicted the degree of behavioral effects of stimulation. Combined, both behavioral and electrophysiological effects of stimulation provide strong causal evidence for the importance of PCC during episodic memory encoding and highlight that memory can be modulated selectively via stimulation of brain structures outside of the hippocampal memory network.

Our key finding is that high-frequency stimulation delivered during encoding resulted in a decrease in observed fraction of successfully recalled items as compared to when there was no stimulation. This result was surprising given that PCC is a critical hub within the default mode network (DMN) (31, 32). Regions of the DMN predominantly show less fMRI activity during encoding of remembered than forgotten items (20–24, 29), which would lead to the plausible hypothesis that stimulating the PCC during encoding would enhance memory. On the contrary, our memory impairment result is consistent with prior reports of negative effects of stimulation in the core memory regions including the temporal lobe and hippocampus (8, 9, 11–13) providing evidence for a causal role of the PCC in memory encoding. Critically, the magnitude of memory disruption during encoding was also correlated with the magnitude of hippocampal-induced gamma band power changes, indicating a potential coupling between PCC and the hippocampal network during encoding. Specifically, stimulation-induced gamma power increases in the hippocampus were associated with increased memory impairment, however, there were substantially reduced differences in gamma power during successful versus unsuccessful item encoding. Thus stimulation produced opposite effects in hippocampal gamma power during successful memory encoding (44, 47–49, 54). One possibility is that stimulation-induced modulations are disruptive to the baseline gamma oscillations in the hippocampus, such that stimulating an already efficiently functioning memory network may negatively impact memory encoding operations. This may further suggest that the PCC exerts causal influence on the genesis of the hippocampal gamma SME routinely observed in memory experiments.

*What is the role of the PCC in memory encoding?* Stimulating the hippocampus and medial temporal lobe leads to a global decline in all facets of episodic memory including memory for primacy items, and temporal and semantic clustering scores (11, 15). This result is consistent with these regions’ critical roles in mnemonic processes and encoding of episodic concepts (55). Our findings of impairment of recall memory, specifically, recollection of primacy items, and possible enhancement of associative memories, reflects a differentiated role for the PCC in supporting episodic memory as compared to the hippocampus or medial temporal lobe. Prior studies speculate that people tend to rehearse early items in a serial position throughout the later list positions (56) and rehearsal provides an encoding boost for early items (49). Thus, one possibility is that PCC plays a role in this rehearsal process and stimulating it during encoding disrupts either the ability to rehearse or impedes benefits of improved representations elicited by rehearsal. Support for this hypothesis is found in fMRI reports showing increase in PCC activation during item rehearsal (57) implicating PCC’s role in creation of robust memory representations via rehearsal mechanisms (58). As such, our data are inconsistent with a role for the PCC in the representation of temporal or semantic associations, but a future exploration of the impact of PCC stimulation in tasks that do not place strong demands on temporal information including associative recognition (59) will help elucidate our observations.

The finding that PCC stimulation affected hippocampal oscillations during memory encoding further bolsters its role in the extended hippocampal memory system (39). Reports of attenuated functional connectivity between the PCC and hippocampus have been associated with mild cognitive impairment and Alzheimer’s disease (38). Consistent with these findings, we observed that the degree of memory impairment is linked to baseline functional connectivity between PCC and hippocampus. We find that participants with larger hippocampal-PCC gamma phase coherence during encoding exhibit greater memory disruption with stimulation, reinforcing the notion that stimulation might be disruptive when delivered to an optimally functioning memory network. Furthermore, our connectivity results highlight the predictive power of functional connectivity and its usefulness in understanding how electrically stimulating a network hub would affect neural responses (52) and behavioral performance. Although, we applied stimulation mainly in the gray matter, future studies can test predictability power using functional connectivity between PCC and hippocampus and if applying stimulation in the white versus gray matter alters the degree of stimulation-induced memory and neural effects (52).

A long-term goal of intracranial research is to explore novel brain targets for potential neuromodulation and to develop strategies for brain-machine interface devices to improve cognitive abilities of patients who suffer from memory and cognitive decline. The PCC was an especially interesting target because (i) it consistently showed decreased activation during successful memory encoding in fMRI studies (20–24), and (ii) using deep brain stimulation to treat movement disorders as a model, we had initially hypothesized that inhibition of the PCC via stimulation during memory encoding could inactivate competing networks and promote successful memory encoding. Although we did not observe memory enhancement with stimulation, a confirmation of PCC’s contribution to memory encoding suggests that the PCC may be a propitious target for closed-loop, responsive stimulation strategies because the impact of PCC stimulation is selective and predictable. Our data is the first to report that extended stimulation epochs can be used to safely modulate hippocampal activity with significant implications on recall performance and that stimulation outside of the temporal lobe can be systematically applied to study networks of episodic memory processing. More generally, our experimental paradigm describes a method to use stereo EEG electrode arrays to elucidate the lesser-known contributions of deep cortical regions towards cognitive processing and can serve as a model for researchers testing alternative strategies for the neuromodulation of memory. Our results also highlight that functional connectivity may be a critical measure for determining whether or not an individual may be a candidate for DBS related therapeutic strategies, that is if stimulation is most likely to improve or disrupt function. This is useful *a-priori* information for developing open-loop (i.e. when stimulation is delivered continuously without response to ongoing neural activity) as well as closed-loop (13, 60) stimulation strategies.

In conclusion, we present novel human data in which we explored the mnemonic functions of the PCC using deep brain stimulation. We (i) demonstrate the feasibility of applying extended periods of electrical stimulation to a deep cortical brain region such as the PCC to modulate memory, (ii) explicate potential causal role for the PCC during memory encoding in hippocampal memory network, and (iii) highlight functional connectivity as an informative measure of stimulation effect for enhancing cognitive functions, with implications for therapeutic advancements in memory diseases.

## Materials and Methods

### Subjects

We collected data from 17 participants (ages 19-63, 9 female) implanted with stereo EEG electrodes targeted to stimulate the posterior cingulate cortex (PCC). Patients were recruited from the epilepsy surgery practice at the University of Texas, Southwestern Medical Center. Only patients who had intracranial electrodes placed within the posterior cingulate and were able to participate in cognitive testing were included in the study. All subjects had normal or corrected-to-normal vision and provided written, informed consent. Protocols were approved by the University of Texas, Southwestern Medical Center Internal Review Board on Human Subjects Research.

### Electrode Localization

Localization of electrodes was performed by pre-surgical planning followed by stereotactic placement of depth electrodes and postoperative fusion of CT images with the pre-operative high-resolution MRI Images (MPRAGE, coronally acquired 1 mm slices) as conducted in our prior studies (40, 61). Depth electrodes were labeled using Freesurfer’s auto-segmentation software (http://surfer.nmr.mgh.harvard.edu/) (62). All electrodes placement was confirmed by expert neuroradiology review as a part of standard clinical practice. For all analysis, we focused on the hippocampal and PCC electrodes (**Fig. 1a**). Across all participants, the total number of hippocampal electrodes was 118, with an average of 6.94±2.4 electrodes per participant and the total number of PCC electrodes was 43, with an average of 2.86±1.68 electrodes per participant. Patients performed a verbal episodic memory task while we applied 25s of continuous high frequency, 100 Hz and 2 mAmps direct brain stimulation to the PCC using the depth electrodes.

### Free recall Experiment

Our goal was to test the effect of stimulation on human memory performance when stimulation was applied during episodic memory encoding. We designed a free recall task in which participants first memorized words from a study list in an encoding phase, following which participants performed a timed math distractor task which lasted 20s, and then participants were asked to verbally recall words from memory in any order, in a retrieval phase. There were a total of 20 study lists, and each list comprised of 10 words. Each word in the study list was presented for 1500 ms, followed by a blank screen for 1000 ms. Stimulation to the PCC was applied during the entire duration of the encoding phase of a study list, for half of the total number of study lists (we refer to these lists as stimulation during encoding (ES) lists), while no stimulation was applied during encoding phase of the remaining word lists (we refer to these lists as no stimulation (NS) lists). As compared to prior studies that applied stimulation during encoding of alternate pairs of study items within each study list (11, 15), in our experiment, stimulation to PCC was delivered throughout the encoding period in a random order to largely avoid concerns of post-stimulation electrophysiological changes impacting oscillatory patterns of nearby items.

To measure baseline functional connectivity between the hippocampus and PCC and to avoid poststimulation related electrophysiological changes and artifacts, 11 of the 17 participants in the previous experiment, performed a second similar free recall task in which no stimulation to PCC was applied during the entire experiment.

### Behavioral analysis of free recall task

To test for behavioral effects of stimulation of PCC on the episodic memory for both during stimulation (ES) versus no stimulation (NS) study lists separately, we measured: (i) percentage of correctly recalled words by measuring the number of correctly recalled words (i.e. only words within a given list without intrusions from prior study lists) across all lists in a condition (NS or ES) and dividing by the total number of words across all lists in a condition (NS or ES), (ii) total number of intrusions during stimulation versus no stimulation lists, (iii) percent recall of items based on their position (1st to 10th separately) in the study list, (iii) primacy effect, evaluating percent recall of primacy item (1^st^ item) versus the percent recall of remaining items, (iv) temporal clustering factor (TCF) which is a measure of subject’s tendency to cluster the recalled items based on their temporal proximity during the study period (45, 46). TCF measures the tendency for recall transitions to occur between items presented at nearby list positions: TCF of 1 indicates perfect temporal contiguity, with the subject only making transitions to temporally adjacent items, whereas a TCF of 0.5 indicates that the subject is making random transitions (45). (v) semantic clustering factor (SCF) which is a distance measure of subject’s tendency to cluster recall items based on their semantic associations (45, 46) (based on a distance matrix provided Latent Semantic Analysis, LSA (63)). TCF and SCF were calculated using the behavioral toolbox developed at the University of Pennsylvania (http://memory.psych.upenn.edu/Behavioral_toolbox).

### Oscillatory power analysis

Intracranial EEG was sampled at 1kHz on a Nihon Kohden 2100 clinical system under a bipolar montage with the most medial white matter contact on individual electrodes as reference. Electrodes exhibiting excessive interictal activity, or recording artifact were excluded from the analysis as part of our standard processing pipeline. We focused our neural data analysis on electrodes in the hippocampus. First, for each subject, per electrode and condition (NS or ES), oscillatory power and phase were extracted (using the Morlet wavelet decomposition, wave number=5) for each study item, in a 1600 msec window following item presentation as in our prior work (40). We employed an automated artifact rejection algorithm to exclude noise and spiking activity (40, 49) (using kurtosis threshold=4). Neural data were sampled at 500Hz. As stimulation was applied at 100Hz, we used an adaptive notch filter to remove stimulation-related noise artifacts. We then compared oscillatory power between ES and NS list items at each log-spaced frequency from 2-64Hz by generating t-statistics at each time-frequency pixel, per electrode. Performing statistical tests within electrodes allowed us to then average data across electrodes using the test statistic, which avoids the need to normalize power values. Across all participants the distribution of t-statistics was compared to zero using an unpaired t-test, resulting in p-values for each time window and frequency, which were converted to z-scores. Significant band clusters across participants were obtained at *p*<0.05, corrected for multiple comparisons. To correlate measures of power analysis with behavioral data, per participant, we averaged t-statistics in frequency-time clusters in theta (2-10Hz), alpha (10-14Hz), beta (15-24 Hz), low gamma (25–40), and high gamma (>40 Hz) bands.

### Subsequent memory effect (SME) analysis

This analysis was conducted to examine how oscillatory power differed during successfully remembered and forgotten items (also known as hippocampal subsequent memory effect (SME (47)), during ES versus NS conditions. For each participant, we measured SMEs, as t-statistics contrasting oscillatory power (in each electrode) during successfully recalled item versus forgotten items, for NS and ES conditions separately and obtained average power across all electrodes. We calculated delta SME as SME in NS minus SME in ES condition, then compared the distribution of differences to zero using unpaired t-tests. Resulting p-values for each time window and frequency, were converted to z-scores (significant frequency band clusters across participants were obtained at *p*<0.05, corrected for multiple comparisons). To correlate measures of SME analysis with behavioral data, per participant, we obtained average delta SME in theta (2-10Hz), alpha (10-14Hz), beta (15-24 Hz), low gamma (25–40), and high gamma (>40 Hz) bands.

### Analysis of baseline functional connectivity between hippocampus and PCC electrodes

Connectivity analysis was conducted on 11 out of 17 participants, who performed a separate free recall task in which no stimulation was applied to the PCC. This provided a “baseline” condition to examine neural connectivity between the hippocampus and PCC. Specifically, using the wavelet-extracted power and phase information, per participant, we computed oscillatory synchrony between the hippocampal electrodes and the posterior cingulate electrodes using phase locking statistics (40, 53). Calculation of oscillatory synchrony was performed with a Rayleigh test applied to the distribution of phase differences at each time-frequency pixel across all items, separately for successfully recalled and forgotten items (64). No baseline was removed from the wavelet-extracted power values for the analysis. Phase locking values (PLV) were calculated via wavelet transform (length 6) after down-sampling to 500 Hz from encoding items (successfully recalled and forgotten items separately), followed by a bootstrap test (200 shuffles) to measure significance level. We randomly selected a subset of subsequently non-recalled items for the synchrony analysis to balance the number of successful and unsuccessful encoding trials while using the Rayleigh test. We extracted a single z-score per electrode pair to quantify the difference in PLS (functional connectivity measurement) and averaged z-score values in theta, alpha, beta, and low gamma, and high gamma bands to quantify connectivity across each frequency band.

### Correlation analyses

All correlations were performed using Pearson’s coefficient.

## Authors’ Contributions

VSN analyzed all patient data and wrote the manuscript; JJ performed data collection and analyzed connectivity data. AB and AA contributed to data collection; MR contributed to discussions on the manuscript and wrote the manuscript. BL performed patient surgeries, implanted and localized electrodes in patients, oversaw all components of the study, and wrote the manuscript.

